# Structure-Based Discovery of a Novel Small-Molecule Inhibitor of Methicillin-Resistant S. aureus

**DOI:** 10.1101/781971

**Authors:** Jie Liu, Lina Kozhaya, Victor J. Torres, Derya Unutmaz, Min Lu

## Abstract

The rapid emergence and dissemination of methicillin-resistant Staphylococcus aureus (MRSA) strains represents a major threat to public health. MRSA elaborates an arsenal of secreted host-damaging virulence factors to mediate pathogenicity and blunt immune defense. Panton-Valentine leukocidin (PVL) and α-toxin are pore-forming cytotoxins of recognized importance in the development of invasive MRSA infection and are thus potential targets for antivirulence therapy. We report the X-ray crystal structures of PVL and α-toxin in their soluble, monomeric and oligomeric, membrane-inserted pore states, in complex with n-tetradecylphosphocholine (C_14_PC). The structures reveal two evolutionarily conserved phosphatidylcholine binding mechanisms and their roles in modulating host cell attachment, oligomer assembly and membrane perforation. Moreover, we demonstrate that the soluble C_14_PC compound protects primary human immune cells in vitro against cytolysis by PVL and α-toxin and hence may serve as the basis for the development of novel antivirulence agents to combat MRSA.

## Introduction

Infection with Staphylococcus aureus can cause severe and devastating illness and is one of the leading causes of death by any infectious agent in the United States (Klevens et al., 2007; Lowy, 1998). S. aureus is notorious for its ability to acquire genetic determinants of antibiotic resistance and virulence that enhance fitness and pathogenicity (Chambers and Deleo, 2009; Otto, 2010). Methicillin-resistant S. aureus (MRSA) now accounts for >60% of S. aureus isolates in US intensive care units, severely restricting antibiotic treatment options (Klevens et al., 2007). MRSA also spreads rapidly among healthy individuals in the community, causing predominantly skin and soft tissue infections and a number of unusually severe clinical syndromes, including necrotizing pneumonia and septic shock (Klevens et al., 2007). Disturbingly, MRSA can live in the biofilm state (Jones et al., 2001; Otto, 2008), and it has long been recognized that biofilms increase resistance to antimicrobial agents and the host immune response (Bowler, 2018). Both vancomycin and daptomycin are key last-line agents for treatment of invasive MRSA infections (Liu et al., 2011). Alarmingly, MRSA strains that are even resistant to vancomycin have emerged recently (Courvalin, 2006; Gardete and Tomasz, 2014). For these reasons, the World Health Organization identifies MRSA as one of six ‘high priority’ pathogens that pose an enormous threat to public health (Willyard, 2017). Therefore, new therapeutics with novel mechanisms of action and that interfere with new targets are desperately needed to combat this high threat pathogen.

USA300 is the most prevalent strain of MRSA in the US and represents a growing threat in both the community and healthcare settings (Diekema et al., 2014). Its heightened incidence and severity have been related to the production of a cocktail of cytolytic pore-forming exotoxins mediating virulence and impairing host immune defenses (Diep et al., 2006; Otto, 2010). The pharmacological targeting of these cytotoxins may be highly effective and lead to a lower selective pressure for resistance than traditional antibiotics. Bipartite leukocidins and single-component α-toxin, secreted by S. aureus as water-soluble, monomeric polypeptides, constitute the α-hemolysin subfamily of β-barrel pore-forming toxins (Gouaux et al., 1997). Five different leukocidins have been described, including Panton-Valentine leukocidin (PVL), leukocidin ED (LukED), two γ-hemolysins (HlgAB and HlgCB) and leukocidin AB (LukAB; also known as LukGH), each of which consists of two distinct polypeptides referred to as the S and F subunits (for a review see Alonzo and Torres, 2014). Their cellular tropism and species specificity are determined by the S subunits LukS-PV, LukE, HlgA, HlgC and LukA (Alonzo et al., 2013; Gauduchon et al., 2001; Spaan et al., 2017). The S and F subunits and α-toxin share a unique modular structure consisting of the amino latch and prestem regions and the β-sandwich and rim domains (see Figure 1A) (Guillet et al., 2004; Nocadello et al., 2016; Olson et al., 1999; Pedelacq et al., 1999; Sugawara et al., 2015). The X-ray crystal structures of the membrane-inserted pore oligomer forms of α-toxin, HlgAB and LukGH and of the membrane surface-bound prepore heterooctamer forms of HlgAB and HlgCB have been determined (Badarau et al., 2015; Song et al., 1996; Yamashita et al., 2014; Yamashita et al., 2011). These structures, and supporting biochemical and genetic data (Sugawara et al., 2015; Valeva et al., 1997; Walker and Bayley, 1995; Yokota and Kamio, 2000), suggest that members of this subfamily share a common mechanism of cytolytic action (for reviews see Dal Peraro and van der Goot, 2016; Kawate and Gouaux, 2003). The cytolytic process begins with the binding of soluble toxin monomers to a cell surface receptor (Berube and Bubeck Wardenburg, 2013; Spaan et al., 2017). The membrane-bound monomers then associate to form a nonlytic, oligomeric prepore. Finally, the translocation of the prestem regions across the membrane results in the bilayer-spanning β-barrel pore structure and consequent membrane permeabilization and cell lysis.

**Figure 1.**
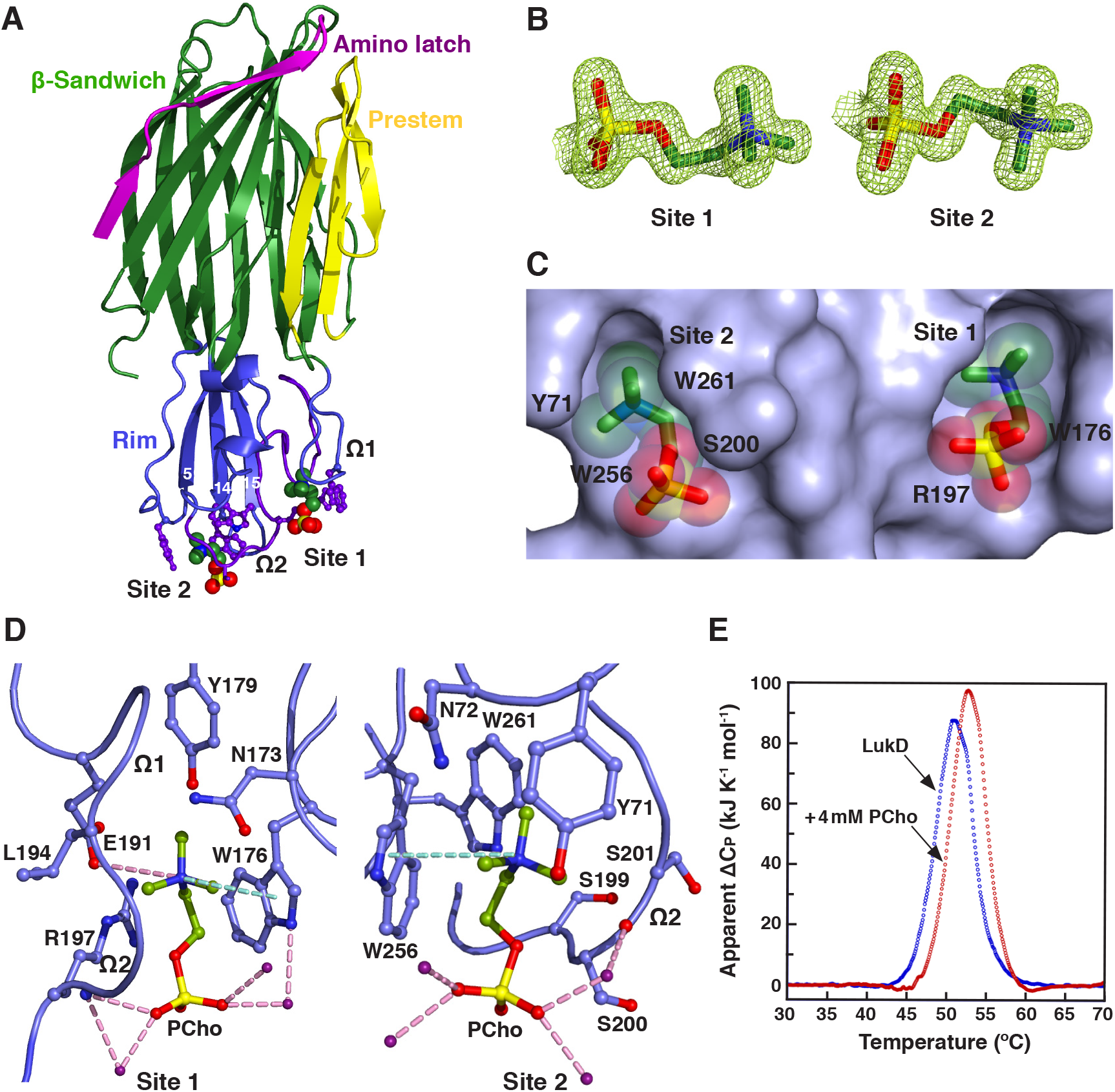
Structural Basis for C_14_PC Binding to LukD. (A) Cocrystal structure of C_14_PC bound to LukD shown from side view perpendicular to the membrane plane, with the PCho moieties and the side chains of key binding site residues displayed as CPK spheres and in ball-and-stick format, respectively. The amino latch (magenta) and prestem (yellow) regions and the β-sandwich (green) and rim (blue) domains are indicated. The two binding sites are labeled, as are the β-strands that compose the rim domain and the Ω1 and Ω2 loops. (B) 2F_o_ – F_c_ omit electron density maps (green mesh) for the two PCho moieties at 1.5σ contour level. (C) Surface representation of the C_14_PC–LukD complex viewed parallel to the membrane, with the PCho moieties shown in stick format with transparent CPK spheres. The locations of key binding site residues are indicated. (D) Close-up view of the two adjacent PCho binding pockets, with residues that make direct side chain contacts with the PCho moieties shown in ball-and-stick format. Cation–π interactions are represented as green dotted lines. Hydrogen bonds and salt bridges are shown as pink dotted lines, and water molecules as purple spheres. (E) DSC thermograms of LukD in the absence (T_m_ = 51.0°C) and presence (T_m_ = 52.8°C) of PCho.

MRSA strains that harbor the phage-encoded PVL have been linked to highly virulent and severe community-acquired skin infections (Lina et al., 1999), as well as life-threatening disease (Gillet et al., 2002). The role of PVL production in the pathogenesis of MRSA was demonstrated in a rabbit model of necrotizing pneumonia (Diep et al., 2010). PVL induces leukocyte destruction and tissue necrosis through interaction with the complement receptors C5aR and C5L2 (Loffler et al., 2010; Spaan et al., 2013; Ward and Turner, 1980; Woodin, 1960). PVL, in conjunction with HlgAB, contributes to MRSA biofilm-mediated killing of neutrophils (Bhattacharya et al., 2018). On the other hand, the chromosomally encoded α-toxin lyses epithelial and endothelial cells, lymphocytes and monocytes by targeting its receptor, the metalloprotease ADAM10 (Berube and Bubeck Wardenburg, 2013; Wilke and Bubeck Wardenburg, 2010). The elevated expression of α-toxin in the USA300 clone and in historic human epidemic strains correlates with increased pathogenicity in mouse models of pneumonia and sepsis (Bubeck Wardenburg and Schneewind, 2008; DeLeo et al., 2011). α-Toxin also plays a role in biofilm formation by clinical MRSA isolates (Anderson et al., 2018). Moreover, LukED relies on the chemokine receptor CCR5 to kill T lymphocytes, macrophages and dendritic cells, as well as CXCR1 and CXCR2 to kill leukocytes (Alonzo et al., 2013; Reyes-Robles et al., 2013). Inhibition of the interaction between LukED and CCR5 has been shown to block cytotoxicity and attenuate S. aureus infection in mice (Alonzo et al., 2013). Together, these cytotoxins can collaborate to modulate phagocytic cell functions via their specific receptors and contribute to MRSA immune evasion and disease pathogenesis. As such, the discovery and development of new antivirulence agents that protect against the combined cytopathic effects of this subfamily of pore-forming toxins are the subject of intense pharmaceutical efforts.

There is considerable evidence pointing to the role of phosphatidylcholine (PC) in the mechanism of pore formation by these toxins. PC is an absolute requirement for pore formation by α-toxin, HlgAB and HlgCB and has been shown to inhibit their cytolytic effects (Ferreras et al., 1998; Noda et al., 1980; Potrich et al., 2009; Valeva et al., 2006; Watanabe et al., 1987). Particularly, crystallographic studies revealed the presence of single, highly conserved phosphocholine (PCho) binding sites on the rim domains of the monomeric F subunit HlgB and the α-toxin protomer in the heptameric pore complex (Galdiero and Gouaux, 2004; Olson et al., 1999). These binding sites have been shown by mutational analysis to be required for membrane targeting and cytolytic function of the two toxins (Monma et al., 2004; Walker and Bayley, 1995). It is generally accepted that the leukocidin F subunits and α-toxin also function in cell attachment through the engagement of their rim domains with the PC head group in the plasma membrane of target cells (Galdiero and Gouaux, 2004; Valeva et al., 2006; Watanabe et al., 1987). In this report, we demonstrate that the soluble, monomeric and oligomeric pore forms of PVL and α-toxin employ two distinct modes to recognize and bind the PC-containing membrane and suggest a novel molecular mechanism for PC-dependent pore formation by members of the α-hemolysin cytotoxin subfamily. Furthermore, we find that n-tetradecylphosphocholine (C_14_PC) effectively inhibits cytolysis of primary human immune cells by PVL, α-toxin and LukED in vitro, thus demonstrating the potential utility of this antivirulence agent alone or in combination with antibiotics against MRSA.

## Results and Discussion

### C_14_PC Binds to the Rim Domain of LukD at Two Adjacent but Distinct Sites

To better understand the molecular basis for the recognition of PCho by the leukocidin F subunits, we determined the crystal structures of LukD with and without C_14_PC at 1.5 Å and 1.75 Å resolution, respectively (Table 1). C_14_PC was selected in the present study as a PC mimic for its high micellization efficiency due to low critical micelle concentration. The two protein structures are closely similar, with a rmsd for Cα atoms of 0.69 Å. The rim domain forms an antiparallel, three-stranded open-face β-sandwich toppled by two surface-exposed consecutive Ω loops (residues 180–194, Ω1 and 195–202, Ω2) (Figure 1A). Two PCho moieties that bind to opposite sides of the Ω2 loop were unexpectedly discovered upon examination of the difference electron density map in the C_14_PC-bound structure (Figure 1B). Average B factor for these two moieties is 22 Å^2^ and for surrounding solvent molecules and protein atoms 21 Å^2^. The two binding sites are approximately 16 Å apart (Figure 1C). The PCho moiety at the first binding site (site 1) is lodged into a concave pocket similar to one in HlgB (PDB code 3LKF). This pocket is formed by two extended segments (residues 171–173 and 176–179, respectively) and the Ω1–Ω2 junction (191– 197) (Figures 1A and 1D). The quaternary ammonium group of the PCho moiety engages in a cation–π interaction with Trp176 while forming a salt bridge to Glu191 (3.79 Å) (Figure 1D). Its N-methyl and methylene groups are in van der Waal contacts (<4.0 Å) with the main chain atoms of Asn173, Glu191, Leu194 and Gly195 and with the side chains of Asn173, Trp176, Tyr179 and Glu191. Furthermore, the phosphate group is hydrogen bonded through its O2 oxygen to the main chain amide of Arg197 (2.85 Å) on one side of the pocket opening, and the side chain of this residue also wraps around the three other oxygens (Figure 1D). In addition, three water molecules form hydrogen bonds to the O2 and O3 oxygens.

**Table 1.**
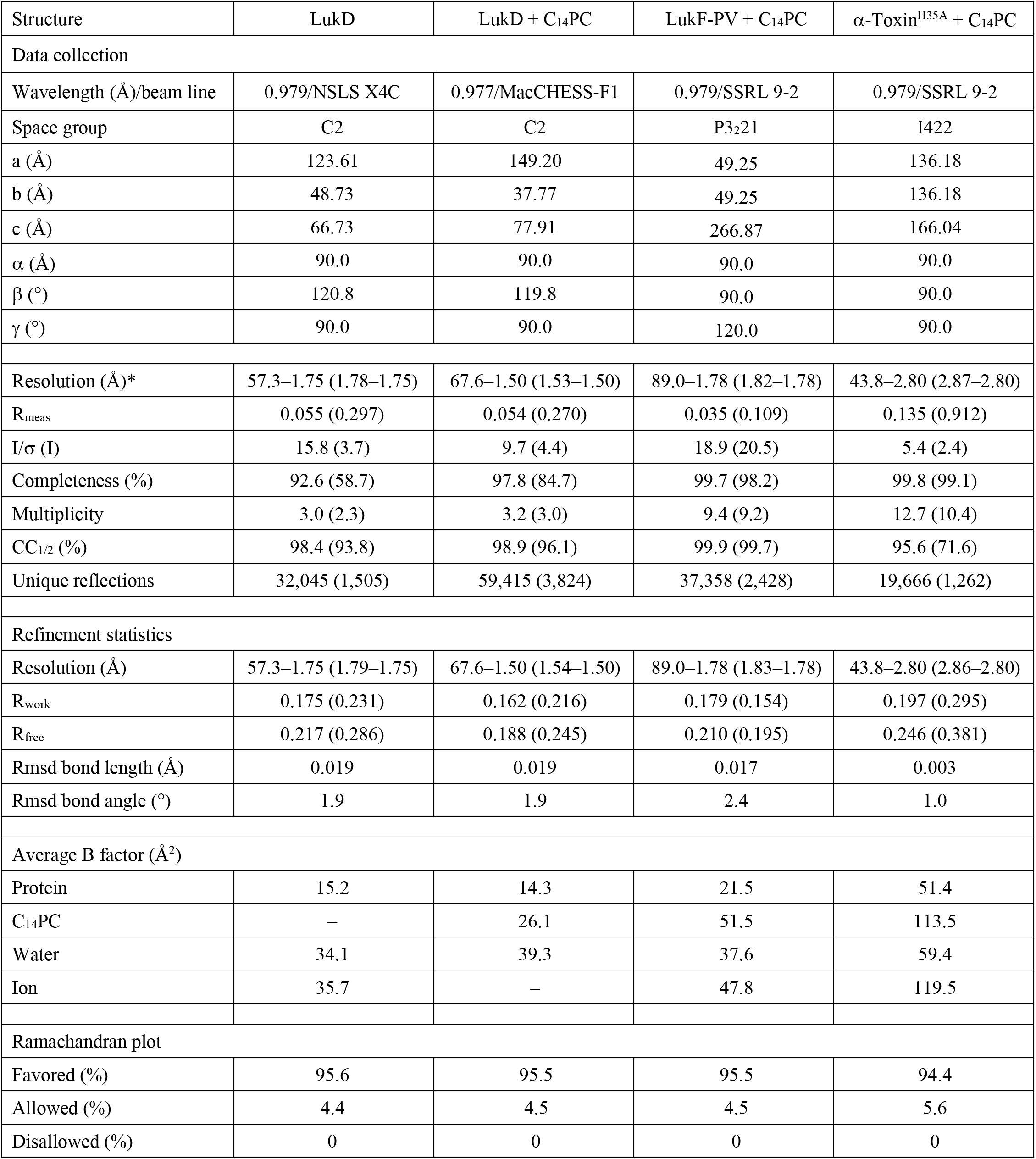
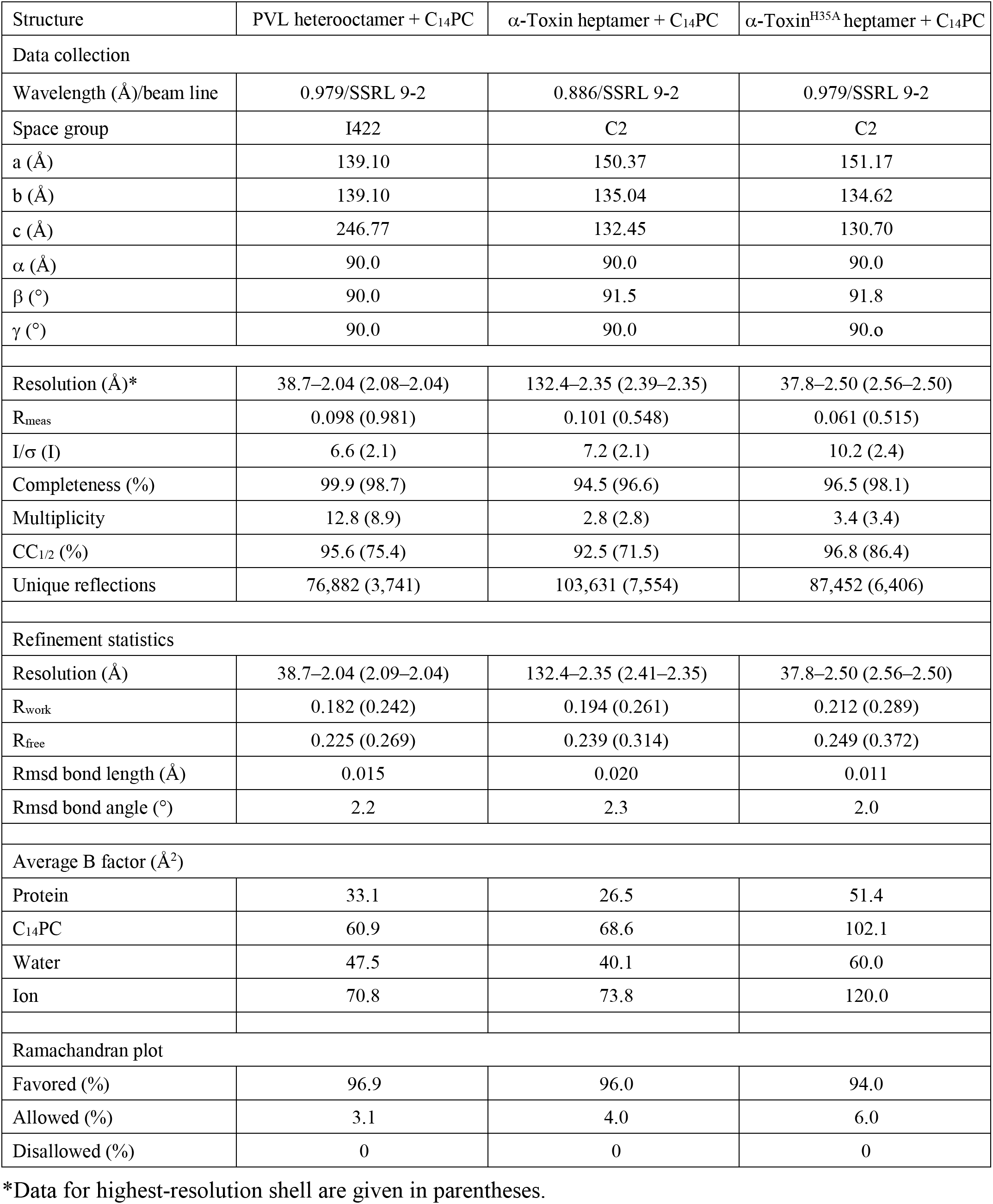
X-Ray Data Collection and Refinement Statistics

| Structure | LukD | LukD + C<sub>14</sub>PC | LukF-PV + C<sub>14</sub>PC | α-Toxin<sup>H35A</sup> + C<sub>14</sub>PC |
| --- | --- | --- | --- | --- |
| Data collection |  |  |  |  |
| Wavelength (Å)/beam line | 0.979/NSLS X4C | 0.977/MacCHESS-F1 | 0.979/SSRL 9-2 | 0.979/SSRL 9-2 |
| Space group | C2 | C2 | P3<sub>2</sub>21 | I422 |
| a (Å) | 123.61 | 149.20 | 49.25 | 136.18 |
| b (Å) | 48.73 | 37.77 | 49.25 | 136.18 |
| c (Å) | 66.73 | 77.91 | 266.87 | 166.04 |
| $\alpha$ (°) | 90.0 | 90.0 | 90.0 | 90.0 |
| $\beta$ (°) | 120.8 | 119.8 | 90.0 | 90.0 |
| $\gamma$ (°) | 90.0 | 90.0 | 120.0 | 90.0 |
| Resolution (Å)* | 57.3–1.75 (1.78–1.75) | 67.6–1.50 (1.53–1.50) | 89.0–1.78 (1.82–1.78) | 43.8–2.80 (2.87–2.80) |
| R<sub>meas</sub> | 0.055 (0.297) | 0.054 (0.270) | 0.035 (0.109) | 0.135 (0.912) |
| I/$\sigma$ (I) | 15.8 (3.7) | 9.7 (4.4) | 18.9 (20.5) | 5.4 (2.4) |
| Completeness (%) | 92.6 (58.7) | 97.8 (84.7) | 99.7 (98.2) | 99.8 (99.1) |
| Multiplicity | 3.0 (2.3) | 3.2 (3.0) | 9.4 (9.2) | 12.7 (10.4) |
| CC<sub>1/2</sub> (%) | 98.4 (93.8) | 98.9 (96.1) | 99.9 (99.7) | 95.6 (71.6) |
| Unique reflections | 32,045 (1,505) | 59,415 (3,824) | 37,358 (2,428) | 19,666 (1,262) |
| Refinement statistics |  |  |  |  |
| Resolution (Å) | 57.3–1.75 (1.79–1.75) | 67.6–1.50 (1.54–1.50) | 89.0–1.78 (1.83–1.78) | 43.8–2.80 (2.86–2.80) |
| R<sub>work</sub> | 0.175 (0.231) | 0.162 (0.216) | 0.179 (0.154) | 0.197 (0.295) |
| R<sub>free</sub> | 0.217 (0.286) | 0.188 (0.245) | 0.210 (0.195) | 0.246 (0.381) |
| Rmsd bond length (Å) | 0.019 | 0.019 | 0.017 | 0.003 |
| Rmsd bond angle (°) | 1.9 | 1.9 | 2.4 | 1.0 |
| Average B factor (Å<sup>2</sup>) |  |  |  |  |
| Protein | 15.2 | 14.3 | 21.5 | 51.4 |
| C<sub>14</sub>PC | – | 26.1 | 51.5 | 113.5 |
| Water | 34.1 | 39.3 | 37.6 | 59.4 |
| Ion | 35.7 | – | 47.8 | 119.5 |
| Ramachandran plot |  |  |  |  |
| Favored (%) | 95.6 | 95.5 | 95.5 | 94.4 |
| Allowed (%) | 4.4 | 4.5 | 4.5 | 5.6 |
| Disallowed (%) | 0 | 0 | 0 | 0 |

**Table 1 (cont.). X-Ray Data Collection and Refinement Statistics**
| Structure | PVL heterooctamer + C<sub>14</sub>PC | α-Toxin heptamer + C<sub>14</sub>PC | α-Toxin<sup>H35A</sup> heptamer + C<sub>14</sub>PC |
| --- | --- | --- | --- |
| Data collection |  |  |  |
| Wavelength (Å)/beam line | 0.979/SSRL 9-2 | 0.886/SSRL 9-2 | 0.979/SSRL 9-2 |
| Space group | I422 | C2 | C2 |
| a (Å) | 139.10 | 150.37 | 151.17 |
| b (Å) | 139.10 | 135.04 | 134.62 |
| c (Å) | 246.77 | 132.45 | 130.70 |
| $\alpha$ (°) | 90.0 | 90.0 | 90.0 |
| $\beta$ (°) | 90.0 | 91.5 | 91.8 |
| $\gamma$ (°) | 90.0 | 90.0 | 90.0 |
| Resolution (Å)* | 38.7–2.04 (2.08–2.04) | 132.4–2.35 (2.39–2.35) | 37.8–2.50 (2.56–2.50) |
| R<sub>meas</sub> | 0.098 (0.981) | 0.101 (0.548) | 0.061 (0.515) |
| I/$\sigma$ (I) | 6.6 (2.1) | 7.2 (2.1) | 10.2 (2.4) |
| Completeness (%) | 99.9 (98.7) | 94.5 (96.6) | 96.5 (98.1) |
| Multiplicity | 12.8 (8.9) | 2.8 (2.8) | 3.4 (3.4) |
| CC<sub>1/2</sub> (%) | 95.6 (75.4) | 92.5 (71.5) | 96.8 (86.4) |
| Unique reflections | 76,882 (3,741) | 103,631 (7,554) | 87,452 (6,406) |
| Refinement statistics |  |  |  |
| Resolution (Å) | 38.7–2.04 (2.09–2.04) | 132.4–2.35 (2.41–2.35) | 37.8–2.50 (2.56–2.50) |
| R<sub>work</sub> | 0.182 (0.242) | 0.194 (0.261) | 0.212 (0.289) |
| R<sub>free</sub> | 0.225 (0.269) | 0.239 (0.314) | 0.249 (0.372) |
| Rmsd bond length (Å) | 0.015 | 0.020 | 0.011 |
| Rmsd bond angle (°) | 2.2 | 2.3 | 2.0 |
| Average B factor (Å<sup>2</sup>) |  |  |  |
| Protein | 33.1 | 26.5 | 51.4 |
| C<sub>14</sub>PC | 60.9 | 68.6 | 102.1 |
| Water | 47.5 | 40.1 | 60.0 |
| Ion | 70.8 | 73.8 | 120.0 |
| Ramachandran plot |  |  |  |
| Favored (%) | 96.9 | 96.0 | 94.0 |
| Allowed (%) | 3.1 | 4.0 | 6.0 |
| Disallowed (%) | 0 | 0 | 0 |

\*Data for highest-resolution shell are given in parentheses.

Immediately adjacent to site 1 is a novel second binding site (site 2), where the PCho moiety occupies a shallow surface pocket that is framed by the C-terminal half of the Ω2 loop (residues 198–202) and the β14–β15 loop (257–260) and flanked by the side chains of Tyr71, Asn72, Trp256 and Trp261 (Figures 1A and 1D). The quaternary ammonium group is sandwiched between the aromatic rings of Tyr71 and Trp256 through cation–π interactions, and the two indole rings of the latter residue and Trp261 interact with each other in an edge-to-face fashion to engage the N-methyl and methylene groups, which also make contacts with the main chain atoms of Ser199, Ser200 and Ser201 and with the side chain of Asn72 (Figure 1D). The phosphate group is secured by a water-mediated hydrogen bonding interaction with the main chain carbonyl of Ser200 (O2–H_2_O = 2.53 Å and H_2_O–O = 2.76 Å), whose Cα and Cα and Cβ atoms pack against the O1, O2 and O4 oxygens (Figure 1D).

The highly complementary interactions between the two adjacent binding sites and the PCho moieties are ostensibly important for specific recognition and binding. The buried solvent accessible surface area of PCho is 262 Å^2^ at site 1 and 231 Å^2^ at site 2, which correspond to approximately 77% and 69% of the unbound PCho surface area, respectively. The side chains of the conserved Trp176–Arg197 and Ser200–Trp256–Trp261 residues, seen below, that define site 1 and site 2, respectively, become more ordered upon binding to C_14_PC. This side chain flexibility could allow these two adjacent, largely preformed pockets to efficiently accommodate the PCho moieties that have distinct binding poses and residue interactions (Figures 1C and 1D). Consistent with this argument, in differential scanning calorimetry (DSC) experiments, LukD (10 μM) was found to unfold in a single cooperative transition, with a midpoint melting temperature (T_m_) of 51.0°C, while this T_m_ value was shifted to 52.8°C in the presence of PCho (4 mM), representing the enhanced thermal stability that accompanies complex formation (Figure 1E). Thus, our results suggest a revised mode of PC recognition and membrane targeting by the rim domain loops.

### Binding Mode of C_14_PC to the Rim Domain of LukF-PV

To validate this binding mode, we cocrystallized LukF-PV with C_14_PC and solved its structure at 1.78 Å resolution (Table 1). In effect, PCho moieties engage the aforementioned two adjacent binding pockets on the rim domain surface (Figures 2A and 2B). At site 1, the quaternary ammonium group of the PCho moiety forms both a cation–π interaction with Trp176 and a salt bridge to Glu191 (3.84 Å); its N-methyl and methylene groups interact with both the main chain atoms of Leu194 and Gly195 and the side chains of Asn173, Trp176, Tyr179, Glu191 and Arg197; and the phosphate group is held in place by a hydrogen bond between its O2 oxygen and the main chain amide of Arg197 (2.72 Å), along with the side chain of this residue lying against the O2 and O3 oxygens (Figure 2B). At site 2, the quaternary ammonium group participates in a cation–π interaction with Trp256 (Figure 2B). Further contacts are made between the N-methyl and methylene groups and both the main chain atoms of Ser199, Asn200 and Leu201 and the side chains of Asn200, Trp256 and Trp261. Polar interactions are also observed between the phosphate and both the main chain atom of Asn200 and the side chain of Asn202 (Figure 2B).

**Figure 2.**
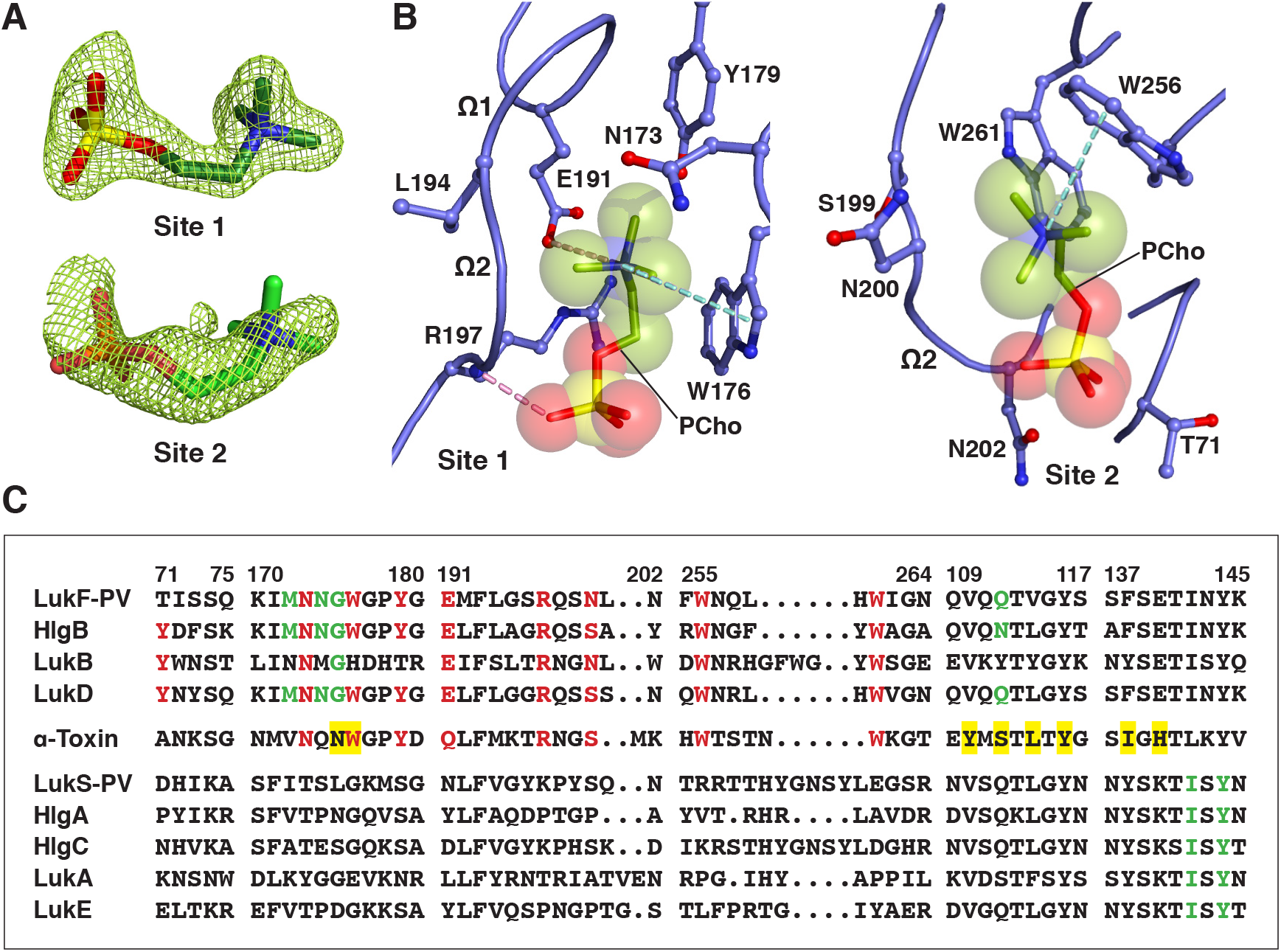
Structure of C_14_PC-Bound Luk-PV. (A) 2F_o_ – F_c_ electron density maps shown as green mesh around the two PCho moieties contoured at 1.5σ (site 1) and 1.0σ (site 2). (B) Molecular interactions in the C_14_PC–LukF-PV complex, with the PCho moieties shown in stick format with transparent CPK spheres. The side chains of residues that make direct contacts with the PCho moieties are shown in ball-and-stick format. The two binding sites and the Ω1 and Ω2 loops are labeled. Cation–π interactions are represented as green dotted lines. Hydrogen bonds and salt bridges are shown as pink dotted lines. (C) Sequence alignment for members of the α-hemolysin pore-forming toxin subfamily around the regions of the three PCho binding sites (see text for details). Four segments of the rim domain (residues 71–75, 170–180, 191–202 and 255–264) and two segments of the stem domain (109– 117 and 137–145) are delineated by spaces and numbered according to the mature LukF-PV protein. Conserved residues at the two binding sites on the rim domain are highlighted red. Conserved residues that constitute the interprotomer binding sites on the PVL heterooctamer and the α-toxin heptamer are highlighted green and a yellow background, respectively.

The solvent accessible surface area of PCho buried by the LukF-PV interaction comprises 264 Å^2^ (79%) at site 1 and 214 Å^2^ (63%) at site 2. DSC measurements reveal that the T_m_ of LukF-PV increased from 50.3°C to 52.3°C when it was bound to PCho. We note that the PCho moiety at site 2 has considerably higher average B factor and poorer electron density than that at site 1 (70 Å^2^ as compared with 31 Å^2^), suggesting that the former moiety is less tightly bound and exhibits greater spatial or temporal disorder. In LukD, the aromatic side chain of Tyr71 contributes to the cation–πbinding interaction to site 2 (see Figure 1D), whereas the corresponding residue in LukF-PV (Thr71) cannot make this interaction (Figure 2C), likely accounting for the lower affinity binding site. The critical functional role of this affinity difference is highlighted by the observation that replacement of Thr71 with a tyrosine endows LukF-PV with the ability to bind human erythrocytes and acquire hemolytic activity when combined with the S subunit of HlgAB (Yokota and Kamio, 2000). Therefore, the elaborate structural features of the two distinct, adjacent PCho binding sites on the leukocidin F subunits may be explained by a selective pressure for membrane PC itself acting as their cell surface receptor.

### C_14_PC Binding by Monomeric α-Toxin^H35A^

To discern the mechanism in the attachment of α-hemolysin subfamily members to host cells, we determined the 2.80 Å crystal structure of C_14_PC in complex with the monomeric His35→Ala mutant of α-toxin (α-toxin^H35A^) (Liang et al., 2009) (Table 1). The asymmetric unit contains two nearly identical protein monomers (rmsd for C_α_ atoms of 0.44 Å), each bound to two PCho moieties (Figure 3A). These moieties occupy the two adjacent binding pockets described above (Figure 3B). At site 1, which is similar to that on the α-toxin protomer in the heptameric pore complex (Galdiero and Gouaux, 2004), the quaternary ammonium group of the PCho moiety makes a cation–π interaction with Trp179 (Figure 3C). Its N-methyl and methylene groups are surrounded by the main chain atoms of Met197 and Lys198 and by the side chains of Asn176, Gln177, Trp179, Tyr182, Gln194, Met197 and Arg200. Importantly, the O2 oxygen of the phosphate group establishes a strong hydrogen bond to the main chain amide of Arg200 (2.64 Å) that also makes side chain contacts with the O2 and O4 oxygens (Figure 3C). At site 2, the quaternary ammonium group forms a cation–π interaction with Trp260, and the N-methyl and methylene groups interact with the main chain atoms of Gly202, Ser203 and Met204 and with the side chains of Ala73, Asn74 and Trp265 (Figure 3B). The phosphate group is clearly visible in the electron density map, although the fine detail of the oxygens is not clear. There are contacts of 3.19 Å between the phosphate and Ser203 and of 3.62 Å between the phosphate and Trp260 (Figure 3C). Upon binding to α-toxin^H35A^, PCho buries 268 Å^2^ (79%) and 203 Å^2^ (61%) of its solvent accessible surface area at site 1 and site 2, respectively. DSC analysis shows that the addition of PCho increased the T_m_ of α-toxin^H35A^ from 50.8°C to 52.4°C. We also observed that the average B factor for the PCho moiety at site 2 is significantly higher than that at site 1 (112 Å^2^ as compared with 75 Å^2^). As discussed in the preceding section, the decreased affinity of site 2 for PCho may arise from the presence of an alanine at position 73 (corresponding to LukD Tyr71) (Figure 2C).

**Figure 3.**
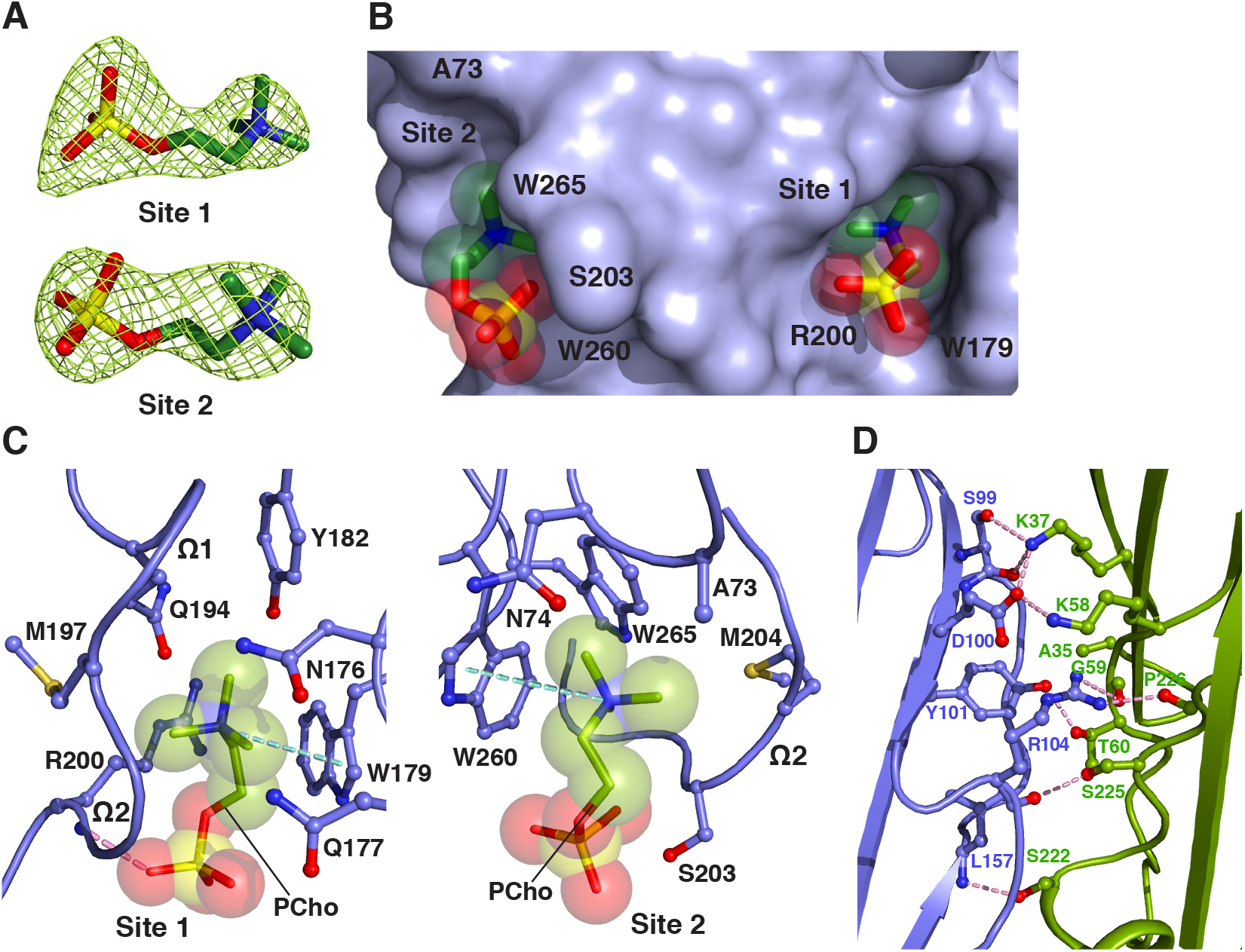
The Two Adjacent PC Binding Pockets on Monomeric α-Toxin^H35A^. (A) 2F_o_ – F_c_ electron density maps (green mesh) for the two PCho moieties contoured at 1.2σ (site 1) and 1.0σ (site 2). (B) Surface representation of the C_14_PC–α-toxin^H35A^ complex viewed parallel to the membrane, with the PCho moieties shown in stick format with transparent CPK spheres. The two binding sites are labeled. The locations of key binding site residues are indicated. (C) Close-up view of the two adjacent PCho binding pockets, with the PCho moieties displayed in stick format with transparent CPK spheres. Residues that make direct side chain contacts with the PCho moieties are shown in ball-and-stick format. The Ω1 and Ω2 loops are labeled. Cation–π interactions are represented as green dotted lines. Hydrogen bonds are shown as pink dotted lines. (D) Close-up view of the interface between the two independent α-toxin^H35A^ monomers (blue and green, respectively) in the asymmetric unit. Hydrogen bonds and salt bridges near the His35→Ala mutation site are depicted as pink dotted lines.

Closer examination of the positions and conformations of the two PCho moieties in the superimposed cocrystal structures of C_14_PC with α-toxin^H35A^, LukD and LukF-PV revealed remarkable similarities. There are few differences in the positions of the five key binding site amino acid side chains (Trp179, Arg200, Ser203, Trp260 and Trp265 in α-toxin; equivalent to Trp176, Arg197, Ser/Asn200, Trp256 and Trp261 in LukD and LukF-PV) in these structures. The three Trp side chains provide two important anchor points for locating the PCho moieties in the two adjacent binding sites, and the Arg and Ser/Asn residues are critical determinants in the binding of the two phosphate groups. Evidently, PC recognition specificity is achieved by a combination of stacking and hydrogen bonding interactions, and van der Waals contacts. Our study shows that membrane PC serves as the common receptor for α-toxin and the leukocidin F subunits, in agreement with previous observations (Galdiero and Gouaux, 2004; Valeva et al., 2006; Watanabe et al., 1987). The presence of the two adjacent PC binding sites on the toxin monomer is consistent with the estimated cross-sectional areas of the PC-bound rim domain (~150 Å^2^) and one PC molecule (~70 Å^2^) (Nagle and Tristram-Nagle, 2000).

Intermolecular contacts between the above two α-toxin^H35A^ monomers comprising the crystal asymmetric unit are formed by residues in the β-sandwich domain (Figure 3D). Comparison of the conformation of these contact residues with their interprotomeric equivalents in the unliganded and C_14_PC-bound heptamers of wild-type α-toxin (PDB code 7AHL; see Figure 5) reveals no local conformational changes involving the main-chain or side-chain atoms. Superposition of the α-toxin^H35A^ dimer onto two adjacent promoters in the above two wild-type toxin heptamers yields overall Cα rmsds of 0.99 and 0.95 Å, respectively, indicating their structural similarity. Dimer interfaces have similar buried surface area values, from 2,061 to 2,171 Å^2^. It is also important to note that the crystal structure of unliganded α-toxin^H35A^ (PDB code 4YHD) lacks the aforementioned intermolecular contacts between six independent monomers in the asymmetric unit. In this structure, both the amino latch and prestem regions have well-defined density with the exception of the six-residue prestem loop and pack against the β-sandwich core of the protein. By contrast, these two regions are apparently disordered in the C_14_PC-bound structure. Our results suggest that α-toxin^H35A^ may be trapped in a PC-bound dimeric state, which may represent an on-pathway intermediate in the assembly of the heptameric pore complex.

**Figure 4.**
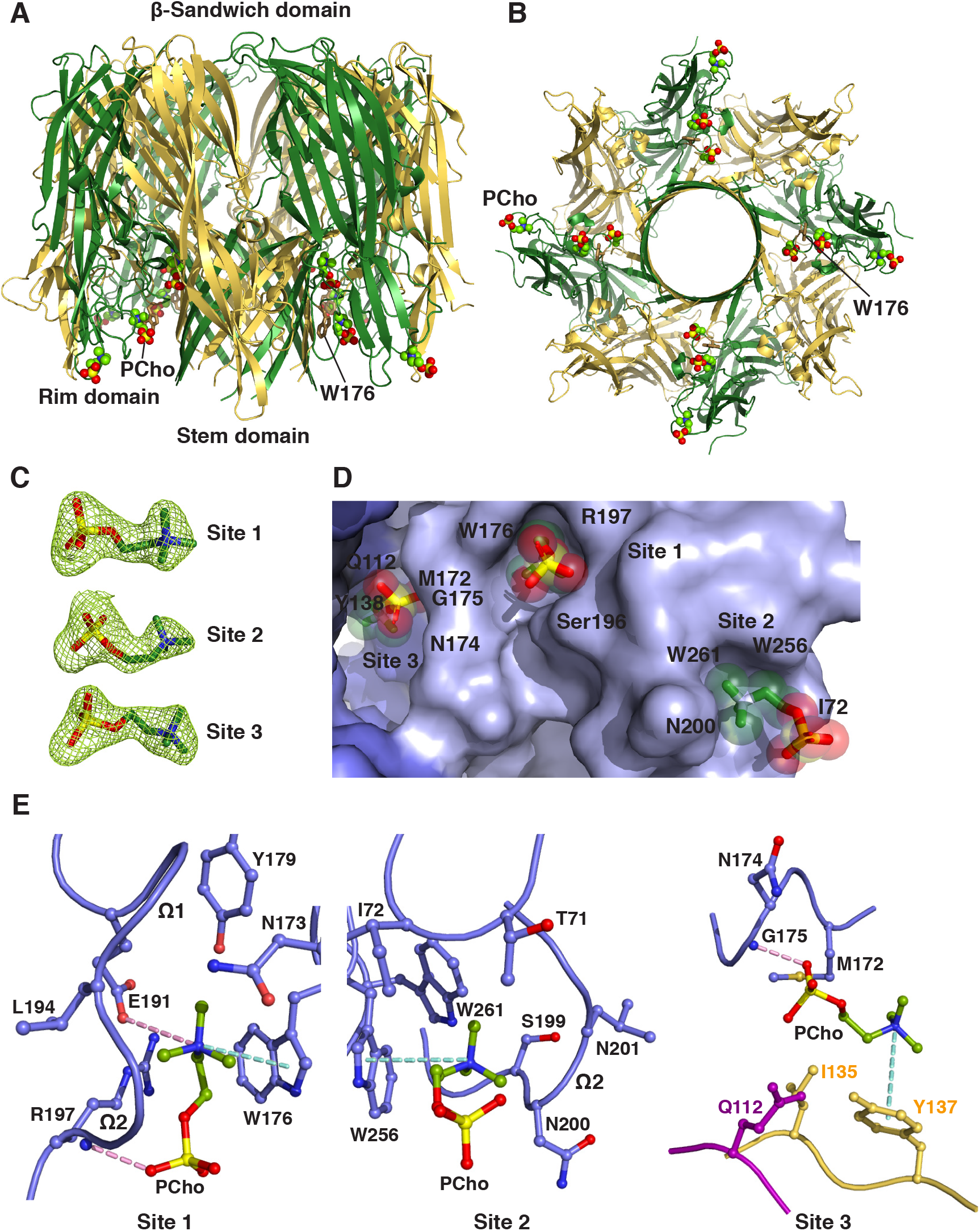
Crystal Structure of C_14_PC in Complex with the PVL Heterooctamer. (A, B) Ribbon representation of the C_14_PC-bound PVL heterooctamer shown from the side (A) and cytoplasmic (B) views, with the PCho moieties and the side chain of Trp176 displayed as CPK spheres and in stick format, respectively. The LukF-PV and LukS-PV subunits are colored green and yellow, respectively. The β-sandwich, rim and stem domains are indicated. (C) 2F_o_ – F_c_ omit electron density maps contoured at 1.0 shown as green mesh around the three PCho moieties in a single protomeric unit. (D) Surface representation of the three PCho binding pockets on a single protomeric unit viewed from the cytoplasmic side, with the PCho moieties shown in stick format with transparent CPK spheres. The three binding sites are labeled. The locations of key binding site residues are indicated. (E) Close-up view of the PCho moieties in the three binding pockets on a single protomeric unit. The side chains of residues that make direct contacts with the PCho moieties are shown in ball-and-stick format colored in blue for protomer A, in magenta for protomer G and in yellow for protomer H. The Ω1 and Ω2 loops are labeled. Cation–π interactions are depicted as green dotted lines. Hydrogen bonds and salt bridges are represented as pink dotted lines.

**Figure 5.**
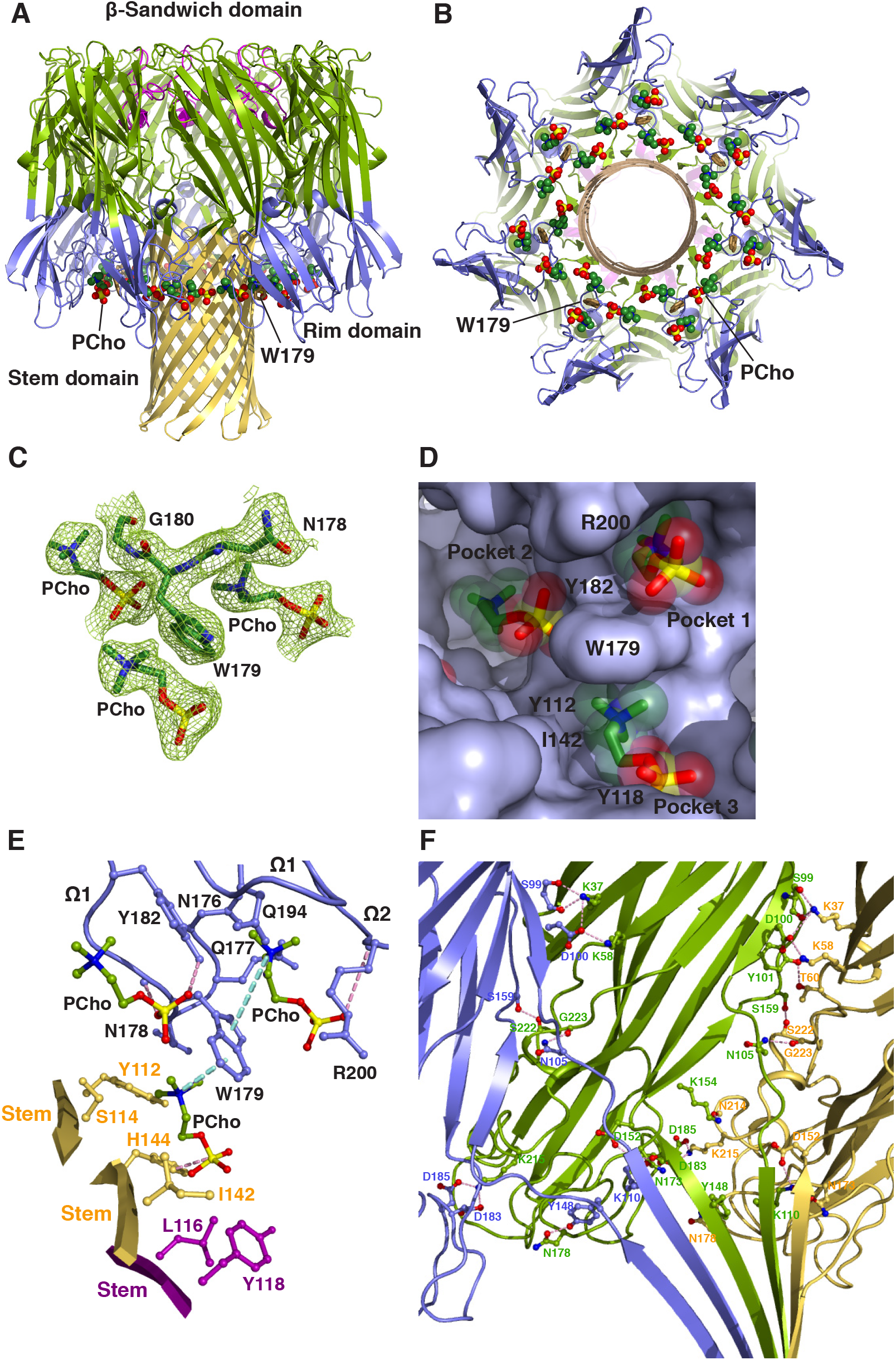
Structure of C_14_PC Bound to the α-Toxin Heptamer. (A, B) Ribbon representation of C_14_PC in complex with α-toxin heptamer shown from the side (A) and cytoplasmic (B) views, with the PCho moieties and the side chain of Trp179 displayed as CPK spheres and in stick format, respectively. The amino latch region and the β-sandwich, rim and stem domains are colored as in Figure 1A. (C) A 2F_o_ – F_c_ electron density map (green mesh) contoured at 1.0σ for residues Asn178, Trp179 and Gly180 and the three PCho moieties in a single protomeric unit. (D) Surface representation of the three partially overlapping PCho binding pockets on a single protomeric unit viewed from the cytoplasmic side, with the PCho moieties shown in stick format with transparent CPK spheres. The three binding pockets are labeled. The locations of key binding site residues are indicated. (E) Molecular interactions in the heptameric α-toxin–C_14_PC complex, with the PCho moieties in a single protomeric unit displayed in ball-to-stick format. Residues that make direct side chain contacts with the PCho moieties are shown in ball-and-stick format colored in blue for protomer A, in purple for protomer E and in yellow for protomer F. The Ω1 and Ω2 loops are labeled. Cation–π interactions are represented as green dotted lines. Hydrogen bonds and salt bridges are shown as pink dotted lines. (F) Close-up view of interprotomer interactions in the triangle region in the crystal structure of the C_14_PC bound α-toxin^H35A^ heptamer. Protomers A, B and C are colored blue, green and yellow, respectively.

Given their expected importance in membrane targeting, the five key PC binding site residues are highly conserved or invariant in both α-toxin and the leukocidin F subunits but are absent in the S subunits, with the exception of a histidine at position 176 in LukB (Figure 2C). Of particular importance, LukB exists as a soluble heterodimeric complex with LukA (DuMont et al., 2014). This finding is consistent with the central role of the conserved Trp176 of the three other F subunits in their binding to the PC bilayer (Olson et al., 1999; Valeva et al., 2006; Watanabe et al., 1987). (and this study). We therefore propose that the binding of the F subunit to the PC-rich membrane is allosterically coupled to heterodimerization with its S subunit counterpart. Likewise, membrane binding by α-toxin, mediated by PC and/or ADAM10, irrevocably commits the monomers to dimerization. The remarkable high degree of conservation of the two adjacent PC binding sites among α-toxin and the F subunits reflects a strong selective pressure on the ability of these two sites to help anchor toxin monomers to the cell surface and to form intermolecular contacts that prime the ensuing formation of the oligomeric, membrane-inserted pore complex.

In summary, the bivalent rim domain interaction with PC provides a mechanism by which soluble toxin monomers can recognize and target the PC-containing membrane, thereby promoting dimer-nucleated pore assembly. The relatively low affinity of PC-mediated binding may facilitate subsequent establishment of the final geometry of the oligomeric pore complex, which we discuss below. α-Toxin and the leukocidin S subunits also bind their specific proteinaceous receptors (Alonzo et al., 2013; Gauduchon et al., 2001; Spaan et al., 2017), and these interactions likely work in concert with the PC targeting mechanism to modulate toxin binding, pore formation and cytotoxicity. Finally, and most importantly, structural elucidation of the two conserved, adjacent PC binding pockets on α-toxin and the leukocidin F subunits will guide the rational development of PC analogs as decoy receptors that effectively divert the cytotoxin away from susceptible cells.

### Structure of the C_14_PC-Bound PVL Heterooctamer

In light of previous studies suggesting that PC plays a crucial role in the assembly and function of the α-toxin heptamer (Galdiero and Gouaux, 2004; Valeva et al., 2006), we cocrystallized the LukS-PV and LukF-PV proteins with C_14_PC in the presence of n-octyl-β-glucoside. The structure of the complex was solved at 2.04 Å resolution by molecular replacement (Table 1). The asymmetric unit contains one LukF-PV/LukS-PV heterodimer and a single LukS-PV molecule. The heterodimer interacts with three crystallographic 4-fold symmetry-related copies of itself to generate a heterooctamer (Figures 4A and 4B). In this β-barrel pore complex, four LukF-PV protomers (denoted A, C, E and G) and four LukS-PV protomers (B, D, F and H) are arranged in an alternating fashion around the central axis of pore, in which the stem domain folds into an antiparallel β-barrel composed of 16 β-strands. We could not discern electron density corresponding to the bottom third of the stem domain in our structure. Two distinct interfaces between neighboring protomers involve residues that are distributed among the amino latch region and the β-sandwich and stem domains, and bury 2,644 Å^2^ and 1,902 Å^2^ of solvent accessible surface area, respectively. The electron density map revealed clearly the presence of PCho moieties at three distinct binding sites on each of the four protomeric units of the PVL heterooctamer (Figures 4C and 4D). The two adjacent binding sites are essentially the same as those on the above-described toxin monomer, whereas the other, novel site lies at the interface between the rim domain of a LukF-PV protomer (e.g., protomer A) and the proximal stem domain regions of protomers G and H. The average B factor for the three PCho moieties is significantly higher than that for the surrounding residues (60 Å^2^ as compared with 31 Å^2^), possibly due to greater disorder and/or subunitary occupancy. Superposition of the PVL hetereooctamer bound to C_14_PC onto the unliganded HlgAB (PDB code 3B07) and LukGH (PDB code 4TW1) heterooctamers yields Cα rmsds of 0.67 and 1.14 Å, respectively, suggesting that the PVL pore does not undergo large conformational changes upon binding to C_14_PC.

The three PCho binding sites on a single protomeric unit are contained within a water-accessible crevice between the inner surface of the rim domain and the upper portion of the stem domain (Figures 4A and 4B). As noted above, the two adjacent sites correspond to those on the rim domain of monomeric LukF-PV (Figure 4D; also see Figure 2B), differing only in the presence of more stabilizing molecular contacts at site 2 on the heterooctamer. Specifically, the quaternary ammonium group of the PCho moiety makes a cation–π interaction with Trp256, and the indole ring of this residue establishes an edge-to-face interaction with the indole ring of Trp261 to pack against the N-methyl and methylene groups, which are also in contact with the main chain atoms of Thr71, Ser199, Asn200 and Leu201 and with the side chain of Ile72 (Figure 4E). At site 3, the aromatic ring of Tyr137 of protomer H forms a cation–π interaction with the quaternary ammonium group and stacks against the N-methyl and methylene groups that are also lined with the side chain of Ile135 of protomer H (Figure 4E). Furthermore, the O3 oxygen of the phosphate group hydrogen bonds to the main chain amide of Gly175 of protomer A (2.65 Å), and the O1 and O3 oxygens engage both the main chain atoms of Asn174 and Gly175 of protomer A and the side chains of Met172 of protomer A and Gln112 of protomer G (Figure 4E). The solvent accessible surface area of PCho buried upon complex formation is 258 Å^2^ (77%) at site 1, 189 Å^2^ (57%) at site 2 and 224 Å^2^ (65%) at site 3.

Our results suggest that multivalent binding of the PVL heterooctamer to PC on the membrane surface leads to localized defects in the lipid bilayer and thus promotes the insertion of amphipathic β-hairpins to produce the β-barrel piercing the bilayer. Critical residues Tyr137 of LukS-PV and Gly175 of LukF-PV at site 3 are invariant in the leukocidin S and F subunits, respectively (Figure 2C), underscoring their functional importance. Furthermore, three similar PC binding pockets also exist in protomers of the C_14_PC-bound α-toxin heptamer described below.

### Binding Mode of C_14_PC to the α-Toxin Heptamer

To evaluate the binding of the α-toxin heptamer to the PC head group in a membrane-mimicking environment, we determined the crystal structure of its complex with C_14_PC at 2.35 Å resolution (Table 1). In this structure, three PCho moieties are bound to each of the seven protomeric units in the water-accessible crevice between the rim and stem domains (Figures 5A and 5B). The indole ring of Trp179 mediates three-way interactions with these three moieties (Figure 5C). Their conformations are clearly defined in three partially overlapping but distinct binding pockets of the crevice (Figure 5D). One pocket corresponds to site 1 on the toxin monomer described above, while the other two are novel heptamer-specific binding sites (see below). The average B factor for the three PCho moieties is 60 Å^2^ and for surrounding protein atoms 33 Å^2^. The structure of the C_14_PC-bound heptamer is very similar to that of the unliganded heptamer (PDB code 7AHL; rmsd for C_α_ atoms of 0.48 Å), with only minor changes in the positions of side chains involved in direct contact with C_14_PC. The pairwise rmsds between protomers A–G in the heptamer span a range from 0.13 to 0.17 Å for C_α_ atoms. The PCho moieties at each of the three binding sites have essentially identical conformation and orientation in each of the seven protomeric units, with average rmsds of 0.34 Å for the first pocket, 0.32 Å for the second pocket and 0.41 Å for the third pocket. For this reason, the following structural analysis of these binding pockets applies to all of the protomeric units.

The first pocket, defined by Trp179 and Arg200, is the same as that on monomeric α-toxin^H35A^ (see Figure 3), albeit the hydrogen bond between the phosphate group of the PCho moiety and the main chain amide of Arg200 is considerably longer and weaker in the latter (Figure 5E). The second pocket lined by all four residues on strand β12 of the rim domain snugly accommodates the PCho moiety (Figures 5D and 5E). It mediates a network of van der Waals contacts involving the main chain atoms of Gly180 and Pro181 and the aromatic rings of Trp179 and Tyr182, forming hydrogen bonds via its hydroxyl group towards the O3 oxygen of the phosphate group (2.69 Å) and via its O2 oxygen with the main chain amide of Gly180 (2.84 Å) while in the cis rotamer.

The third pocket is located at the interface between the rim domain of protomer A and the proximal stem domain regions of protomers E and F (Figure 5E), in contrast to the other pockets that are constituted solely by residues from the rim domain. The third pocket is formed by residues Asn178 and Trp179 from the rim domain of protomer A, by Leu116 and Tyr118 from the stem domain of protomer E and by Tyr112, Ser114, Ile142, Gly143 and His144 from the stem domain of protomer F (Figure 5E). The indole ring of Trp179 is situated to produce a cation–π interaction with the quaternary ammonium group of the PCho moiety (Figure 5E). The N-methyl and methylene groups participate in extensive contacts with the main chain atoms of Gly143 and Asn178 and with the side chains of Tyr112, Ser114 and Ile142. The PCho moiety is further stabilized by a hydrogen bond between the O3 oxygen of the phosphate group and the ND1 atom of His144 (2.78 Å) and by contacts between the O1, O2 and O3 oxygens and the side chains of Leu116, Tyr118 and His144 (Figure 5E). The solvent accessible surface areas buried upon binding of the PCho moieties to the first, second and third pockets are 260 A^2^ (76%), 207 A^2^ (60%), 285 A^2^ (83%), respectively.

These results strengthen the hypothesis that multivalent binding of the PC bilayer by the α-toxin heptamer may help overcome the energetic barrier to deformation of the membrane during assembly of the β-barrel pore lining, thereby driving the prepore-to-pore conversion. Indeed, replacement of Trp179 and Arg200 with alanines in α-toxin is known to lead to an arrested prepore state in which only the top half of the cytolytic β-barrel pore has formed (Sugawara et al., 2015). Together with analysis of intermediate stages of the α-toxin assembly process with engineered disulfide bonds (Kawate and Gouaux, 2003), our study also suggests that the interaction between the α-toxin prepore and the PC head group may induce a large conformational change in the prestem region, which is essential for pore formation.

### Structure of the α-Toxin^H35A^ Heptamer in Complex with C_14_PC

In the α-toxin pore structure, His35 is located in the crucial interprotomeric contact region (Song et al., 1996), and nonconservative replacements at this position (including H35A) have been shown to abolish heptamer formation and thus cytolytic activity and lethal toxicity (Jursch et al., 1994; Krishnasastry et al., 1994; Menzies and Kernodle, 1994). In light of our findings that the PC bilayer binding might promote both the oligomerization of α-toxin monomers and the structural rearrangements that accompany the prepore-to-pore conversion, we hypothesized that a high concentration of C_14_PC could facilitate the assembly of the α-toxin_H35A_ pore complex. To directly test this hypothesis, we have determined the structure of the α-toxin^H35A^ heptamer crystallized in the presence of 25 mM C_14_PC (see Experimental Procedures) at 2.5 Å resolution (Table 1 and Figure 5F). In this mutant pore complex, PCho moieties bind in the first and second pockets described above on the rim domain of each protomer. In essence, the C_14_PC-bound structures of the α-toxin^H35A^ and wild-type heptamers are nearly identical, with rmsds of 0.04–1.42 Å over 2,051 Cα atoms. The positions and conformations of the two PCho moieties are also similar. However, C_14_PC does not bind to the aforementioned interprotomer pocket on the α-toxin^H35A^ pore, while B factors for this mutant pore are considerably higher than those for the wild-type one (24–201 Å^2^ as compared with 13–73 Å^2^), consistent with the pronounced effect of the H35A mutation on cytotoxicity (Liang et al., 2009). These results support our hypothesis that the PC-rich membrane acts as a critical effector of oligomerization and pore formation by α-toxin.

In summary, despite their different subunit composition and stoichiometry, α-toxin and the leukocidins likely follow an evolutionarily conserved PC-dependent pore assembly pathway, involving the initial membrane binding of toxin monomers, dimerization and oligomerization, and the prepore-to-pore transition and membrane perforation. Importantly, atomic-level insight of the toxin oligomer–PC interactions obtained here will facilitate the development of PC analogs that antagonize pore formation and thus block the cytolytic effects of this subfamily of proteins.

### Inhibition of the Cytotoxicity of LukED, PVL and α-Toxin by C_14_PC

The presence of the conserved PC binding sites in the leukocidins and α-toxin (see above) suggests that PC mimetic compounds may confer protection from toxin-mediated killing of primary human immune cells. Therefore, flow cytometry experiments were conducted to first evaluate the ability of C_14_PC to diminish the cytolytic activity of LukED in Jurkat cells expressing CCR5. This Jurkat cell line has been shown to be susceptible to the toxin (Alonzo et al., 2013). LukED at a concentration of 2.5 μg/mL resulted in ~80% killing of Jurkat cells within 1 h at 37°C (Figure 6A). We found that C_14_PC inhibited the lysis in a concentration-dependent manner, with IC_50_ values of 15–25 μM (Figure 6A). In sharp contrast, PCho does not have appreciable inhibitory activity. We conclude that C_14_PC produces effective toxin inhibition by presenting multiple copies of the PC head group on its micellar surface, in accordance with previous observations (Valeva et al., 2006).

To determine the capacity of C_14_PC to abrogate LukED cytotoxicity toward primary human leukocytes expressing CCR5 and CXCR1 chemokine receptors in vitro, LukED at concentrations of 2.5 and 5 μg/ml was first preincubated with 50 μM C_14_PC at 4°C, and was subsequently added to PBMCs labelled with specific cell surface markers. After 1–1.5 h at 37°C, the cells were stained with fixable viability dye eFluor 506 and analyzed by flow cytometry. Inhibition of LukED by C_14_PC was assessed by determining the relative abundance of viable cells after challenge with the toxin or media. As expected, CD14^+^ monocytes were significantly absent by 2.5 and 5 μg/mL of LukED (Figure 6D), while pretreatment with 50 μM C_14_PC produced a 70–90% protective effect against monocyte lysis (Figures 6B and 6D). Likewise, 50 μM C_14_PC efficiently blocked the lysis of CD8^+^ effector memory T cells by 50–75% (Figures 6C and 6E) and CD8^+^CCR5^+^ T cells by 50– 95% (Figure 6F). Moreover, inhibition of the cytolytic action of LukED by 50 μM C_14_PC also rescued 50–85% of NK cells (Figure 6G), which are highly susceptible to the toxin due to their surface expression of CXCR1 (Alonzo et al., 2013). These results demonstrate that C_14_PC can confer broad-spectrum protection against LukED-mediated killing of target host cells by virtue of its inhibitory effect on the interaction between the toxin and membrane PC.

**Figure 6.**
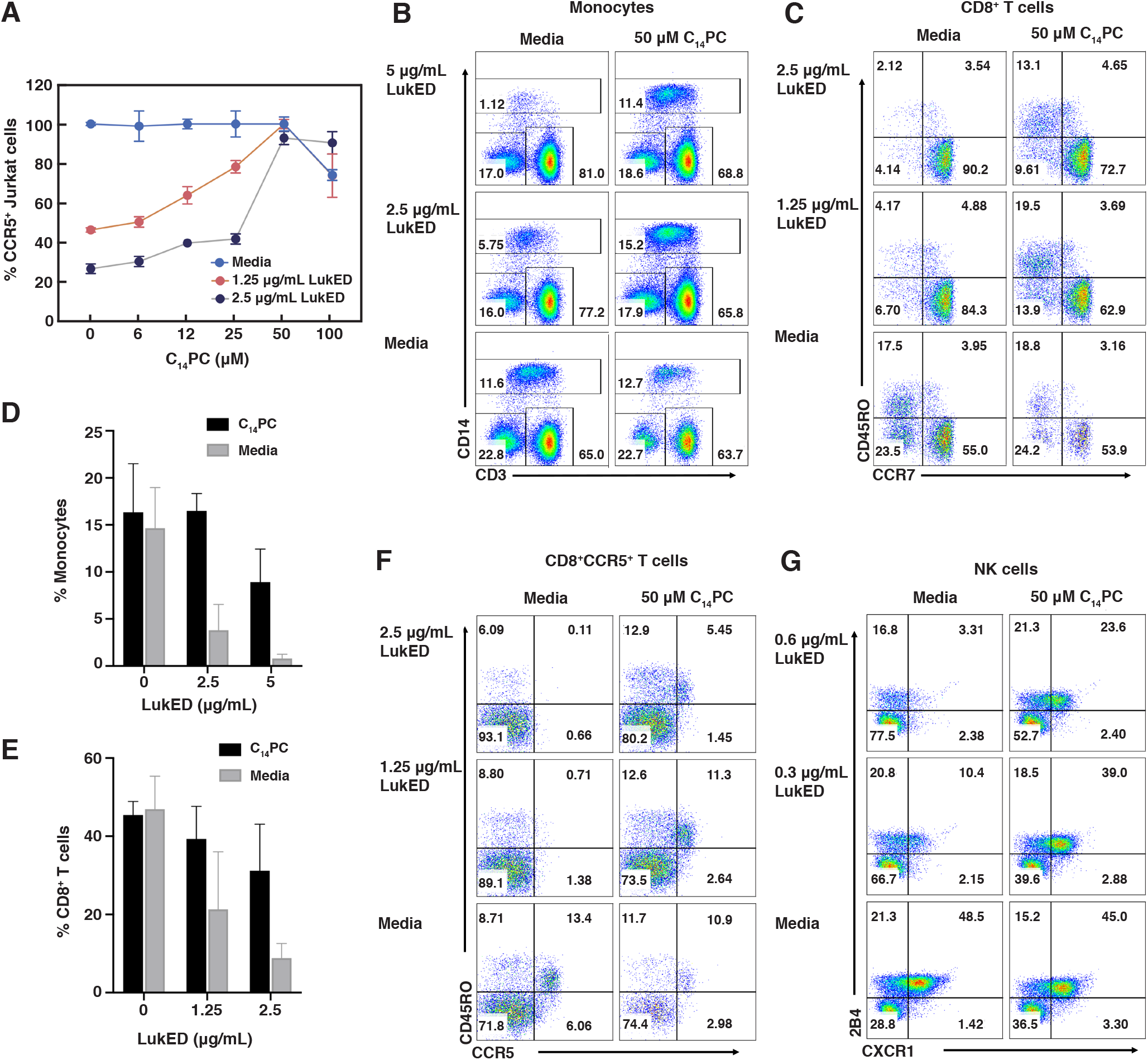
C_14_PC Protects Target Cells from LukED-Mediated Cytolysis. (A) Titration of LukED cytotoxicity by C_14_PC. CCR5^+^ Jurkat cells were challenged with different concentrations of LukED in the presence and absence of C_14_PC. Cell viability was determined by flow cytometry. Data are representative of at least three independent experiments and values are expressed in the mean of triplicate measurements ± standard error (SE). (B) Protective effect of C_14_PC against LukED killing of primary human monocytes. PBMCs were challenged with LukED in the presence and absence of C_14_PC. Monocyte subsets were identified as CD3^−^CD14^+^ after gating on live PBMCs as determined by flow cytometry. Data are representative of two independent experiments. Percentages of cells in each quadrant are indicated. (C) Protective effect of C_14_PC against LukED killing of CD8^+^ effector memory T cells (CCR7^−^CD45RO^+^ and CCR7^−^CD45RO^−^). PBMCs were challenged with LukED in the presence and absence of C_14_PC. Live CD3^+^CD8^+^ T cell subsets were gated and analyzed for the expression of CCR7 and CD45RO by flow cytometry. Data are representative of two independent experiments using blood from different donors. (D) Bar graph showing the inhibition of LukED-mediated cytolysis of monocytes after pretreatment with C_14_PC. Error bars indicate SEM. (E) Bar graph showing C_14_PC inhibition of LukED-induced lysis of CD8^+^ effector memory T cells. Error bars indicate SEM. (F) Protective effect of C_14_PC against LukED killing of CD8^+^CCR5^+^ T cells. PBMCs were challenged with LukED in the presence and absence of C_14_PC. Live CD3^+^CD8^+^ T cell subsets were gated and analyzed for the expression of CCR5 and CD45RO by flow cytometry. Data are representative of two independent experiments. (G) Protective effect of C_14_PC against LukED killing of NK cells. PBMCs were challenged with LukED in the presence and absence of C_14_PC. PBMCs were first gated on live CD3^−^HLA^−^DR^−^ cells and proportion of CXCR1^+^2B4^+^ NK cells was analyzed by flow cytometry. Data are representative of two independent experiments.

We next sought to assess the ability of C_14_PC to inhibit the cytolytic activities of PVL and α-toxin using the in vitro cell viability assay described above. Addition of 10 ng/mL of PVL led to nearly complete lysis of monocytes after 1.5 h of incubation at 37°C (Figure 7A). Pretreatment with 100 μM C_14_PC suppressed the lysis by 90% (Figures 7A and 7B). Similarly, α-toxin at concentrations of 30 and 100 ng/mL caused 75–90% lysis of monocytes and ~50% lysis of CD3^+^ T cells after incubation at 37°C for 24 h, while 100 μM C_14_PC reduced the cytotoxic activity of the toxin by 75–90% (Figures 7C and 7D). We conclude that C_14_PC is a broad-spectrum small-molecule inhibitor of the α-hemolysin subfamily of toxins and that membrane PC contributes to the mechanism of their cytolytic action.

**Figure 7.**
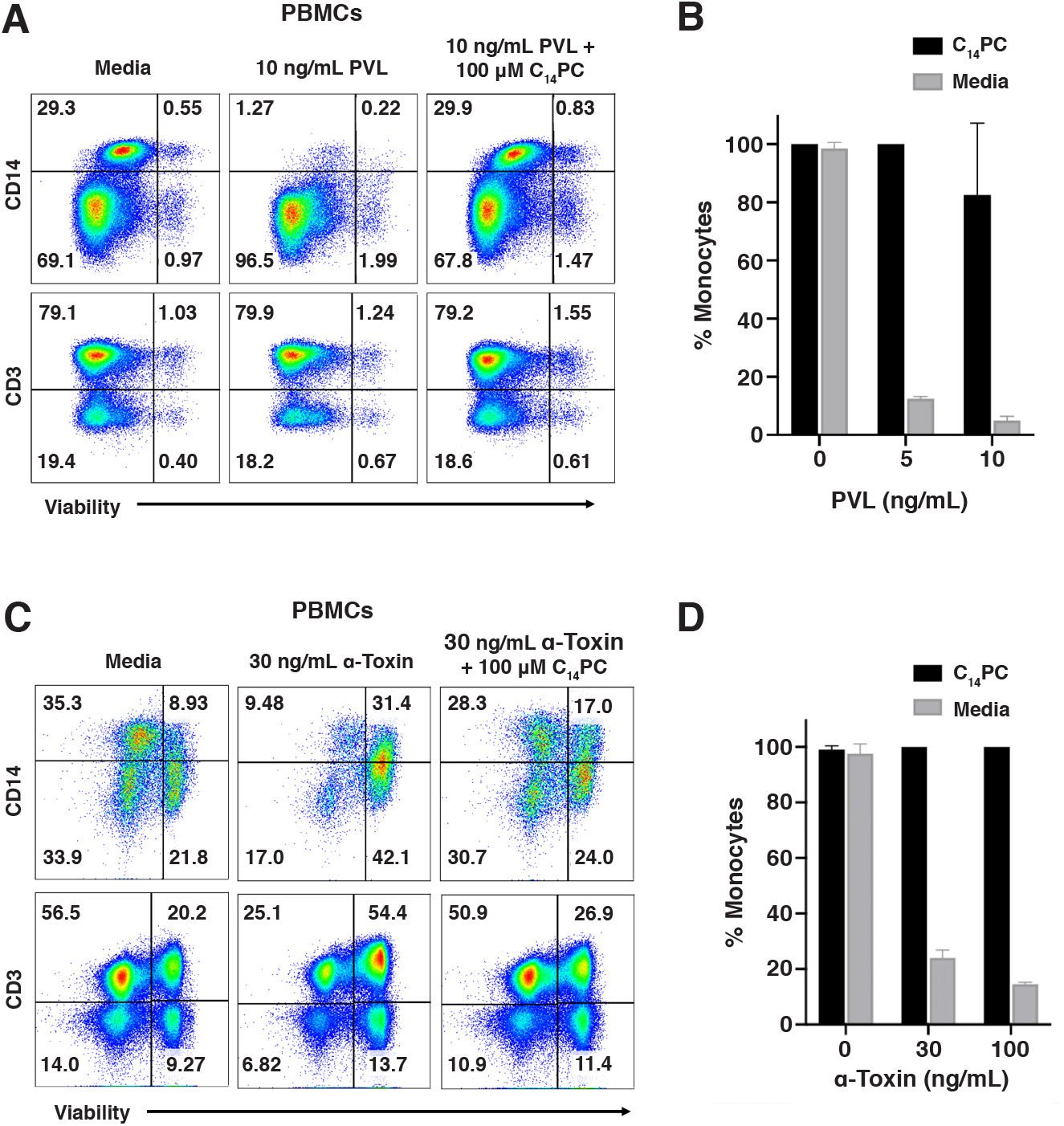
Protection of Primary Human Monocytes Against PVL and α-Toxin by C_14_PC. (A) Protective effect of C_14_PC against PVL killing of monocytes. PBMCs were challenged with PVL in the presence and absence of C_14_PC. Monocyte subsets were gated based on the FSC and SSC parameters and then analyzed for the expression of CD14^+^ and CD3^+^, respectively. Cell viability was determined by flow cytometry and normalized to that in the medium control. Data are representative of two independent experiments using blood from different donors. Percentages of cells in each quadrant are indicated. (B) Bar graph showing the protection from PVL-mediated cytolysis of monocytes by C_14_PC. Error bars indicate SEM. (C) Protective effect of C_14_PC against α-toxin-mediated killing of monocytes. PBMCs were challenged with α-toxin in the presence and absence of C_14_PC. Monocyte subsets were identified and quantified as described in A. Data are representative of two independent experiments using blood from different donors. (D) Bar graph showing the inhibition of α-toxin-induced lysis of monocytes by C_14_PC. Error bars indicate SEM.

### Implication for MRSA Drug Discovery

The high prevalence of multidrug-resistant S. aureus and its remarkable pathogenic potential are creating a crisis in modern healthcare due to the limited therapeutic options available, the toll of severe disease and mortality it inflicts, and the enormous cost of inpatient care to which it contributes (Chambers and Deleo, 2009; Otto, 2010). The ability of MRSA to form biofilms on necrotic tissues and medical devices is also an important virulence mechanism that complicates infections (Jones et al., 2001; Otto, 2008). As antibiotics are becoming less effective, disarming the major virulence mechanisms of MRSA strains has potential to become an alternative, viable therapeutic approach aimed at limiting host tissue damage while aiding immune clearance. The α-hemolysin subfamily of cytotoxins are prime targets for novel therapeutics, owing to their critical roles in inactivating host immune defenses, destroying tissue barriers and modulating inflammatory responses (Diep et al., 2006; Otto, 2010). Currently, neutralizing antibodies targeting α-toxin are in clinical trials (Hua et al., 2015; Le et al., 2016). Given the variability of MRSA immune evasion determinants, such single-target drugs are most likely to be inadequate to achieve a therapeutic effect (Sause et al., 2016). Our structural elucidation of the conserved role of membrane PC in pore formation by members of this subfamily offers new opportunities for concurrent intervention of their pathology via blocking both toxin binding to host cells and pore complex assembly. As a result of our combined structural biology and pharmacological approach, we have been able to demonstrate that C_14_PC is a novel broad-spectrum inhibitor of the leukocidins and α-toxin in vitro. In light of the safety of miltefosine (hexadecylophosphocline, C_16_PC), an oral drug used for the treatment of leishmaniasis (Dorlo et al., 2012), it is reasonable to surmise that the C_14_PC will likewise be well tolerated in humans. Considering its multi-target mechanism of action and low production costs, C_14_PC could potentially be developed as a prophylactic and therapeutic agent against MRSA.

## Experimental Procedures

### Chemicals

All chemicals used were of analytical grade. Unless otherwise indicated, chemicals were purchased from Sigma-Aldrich. Detergents were from Anatrace.

### Cloning and Protein Purification

The full-length LukD (residues 1–301), LukE (1–283), LukF-PV (1–301), LukS-PV (1–284) and α-toxin (1–293) constructs, excluding their signal peptides, were subcloned individually into a modified pET3a vector (Novagen). Site-directed mutagenesis was carried out using the Kunkel method. All constructs were verified by DNA sequencing. E. coli BL21(DE3)pLysS cells transformed with each plasmid were grown at 37°C in LB medium until the A_600_ was between 0.6 and 0.8. IPTG was then added to a final concentration of 0.5 mM, and incubation was continued for 24 h at 16°C. Cells were harvested by centrifugation, suspended in 50 mM sodium acetate (pH 5.4), 25% sucrose, 5 mM EDTA and 5 mM DTT and lyzed at 4°C using an Avestin Emulsiflex C3 homogenizer. Inclusion bodies were isolated by centrifugation, washed twice with the same buffer and subsequently incubated overnight at 4°C in 50 mM sodium acetate (pH 5.4), 5 mM DTT and 6 M guanidine-HCl or 8 M urea. Insoluble material was removed by centrifugation, and the protein solution was then dialyzed for 2 days at 4°C against three changes of buffer A (50 mM sodium acetate [pH 5.4] and 1 mM EDTA). After removal of the insoluble material by centrifugation, the refolded recombinant toxin was loaded onto a CM-Sepharose CL-6B column equilibrated with buffer A and eluted using a linear gradient from 0 to 1 M NaCl. Fractions containing the toxin were pooled, dialyzed against buffer A, concentrated and loaded onto a GE Mono S 5/50 GL equilibrated with buffer A, and the toxin was eluted using a linear gradient from 0 to 0.5 M NaCl. The toxin was further purified using size exclusion chromatography on a GE Superdex 200 10/300 GL equilibrated with 50 mM sodium acetate (pH 5.4) containing 100 mM NaCl. Fractions containing the toxin were pooled, concentrated to ~20 mg/mL and stored at –80°C until use. The concentration of the toxin in purified preparations was determined through UV absorbance measurements.

### Crystallization

All crystallization experiments were performed at room temperature using the hanging drop-vapor diffusion method by mixing 1 μL of protein solution with an equal volume of reservoir solution. Crystals of LukD were grown from a protein solution (12 mg/mL) in 10 mM sodium acetate (pH 5.4) and a reservoir solution containing 20% PEG MME 2000, 10 mM NiCl_2_, 0.1 M Tris-HCl (pH 8.5). For data collection, the crystals were cryoprotected with 15% glycerol in the mother liquor and then flash-cooled in liquid nitrogen. The C_14_PC–LukD complex was crystallized from a protein solution (10 mg/mL) in 10 mM sodium acetate (pH 5.4), 10 mM C_14_PC, 30 mM n-octyl-β-D-glucoside (βOG) and a reservoir solution containing 28% PEG 400, 0.2 M MgCl_2_, 0.1 M HEPES (pH 7.5). The crystals were flash-cooled by plunging directly into liquid nitrogen. Crystals of LukF-PV complexed with C_14_PC were grown from a protein solution (10 mg/mL) in 10 mM sodium acetate (pH 5.4), 10 mM C_14_PC, 30 mM βOG and a reservoir solution containing 2.6 M ammonium sulfate, 5% PEG 400, 0.1 M HEPES (pH 8.5). The crystals were flash-cooled in liquid nitrogen. The C_14_PC–α-toxin^H35A^ complex was crystallized from a protein solution (10 mg/mL) in 10 mM sodium acetate (pH 5.4), 5 mM C_14_PC, 40 mM βOG, 0.4 mM Deoxy-Big CHAP and a reservoir solution containing 1.5 M ammonium sulfate, 0.25 M potassium sodium tartrate, 0.1 M sodium citrate (pH 6.0). The crystals were transferred into a stabilizing solution containing 2.25 M ammonium sulfate, 5% glycerol, 20 mM C_14_PC, 0.1 M sodium citrate (pH 6.0) and then allowed to equilibrate against 3 M ammonium sulfate for 1 h at room temperature prior to flash freezing in liquid nitrogen. The PVL heterooctamer in complex with C_14_PC was crystallized from a protein solution (6.7 mg LukF-PV/mL and 6.3 mg LukS-PV/mL) in 10 mM sodium acetate (pH 5.4), 15 mM C_14_PC, 40 mM βOG and a reservoir solution containing 0.16 M magnesium formate. The crystals were transferred into a dehydrating solution containing 2.7 M ammonium sulfate, 20 mM C_14_PC and then allowed to equilibrate against 3 M ammonium sulfate for 3 h at room temperature prior to flash freezing in liquid nitrogen. Crystals of the α-toxin heptamer–C_14_PC complex were grown from a protein solution (8 mg/mL) in 10 mM sodium acetate (pH 5.4), 15 mM C_14_PC, 30 mM βOG and a reservoir solution containing 2 M ammonium sulfate, 0.2 M potassium sodium tartrate, 0.1 M sodium citrate (pH 6.0). The crystals were flash-frozen in liquid nitrogen. The α-toxin^H35A^ heptamer in complex with C_14_PC was crystallized from a protein solution (10 mg/mL) in 10 mM sodium acetate (pH 5.4), 25 mM C_14_PC, 40 mM βOG and a reservoir solution containing 1.9 M ammonium sulfate, 0.25 M potassium sodium tartrate, 0.1 M sodium citrate (pH 5.2). The crystals were flash-cooled in liquid nitrogen.

### X-Ray Data Processing and Crystallographic Refinement

Diffraction data were collected at 100 K at beamline X4C at the National Synchrotron Light Source at Brookhaven National Laboratory, at the Cornell High Energy Synchrotron Source (CHESS) beamline F1 and at the Stanford Synchrotron Radiation Lightsource (SSRL) beamline 9-2. The diffraction data were processed with HKL-2000 (Otwinowski and Minor, 1997). Initial phases were determined by molecular replacement using Phaser (McCoy et al., 2007) with respective models of HlgB (PDB code 1LKF), LukF-PV (1PVL), α-toxin^H35A^ (4YHD), the HlgAB heterooctamer (3B07) and the α-toxin heptamer (7AHL). Refinement was carried out in Refmac5 (Murshudov et al., 1997), alternating with manual rebuilding and adjustment in COOT (Emsley and Cowtan, 2004). Coordinates for the C_14_PC molecule were generated using LibCheck (Vagin et al., 2004). TLS refinement was performed in Refmac5 (Winn et al., 2001). Detailed collection and refinement statistics are summarized in Table 1.

### Structural Analyses

Model quality was judged using the programs Rampage, Procheck and Sfcheck (Laskowski et al., 1993; Lovell et al., 2003; Vaguine et al., 1999). Protein-ligand contacts for the toxin–C_14_PC complex structures were analyzed using the program COOT (Emsley et al., 2010). The rmsd values were calculated using the program SuperPose (Maiti et al., 2004). Molecular and solvent-accessible surfaces were calculated with the AREAIMOL program (Lee and Richards, 1971) from the CCP4 suite (Winn et al., 2011). PyMOL (DeLano Scientific) was used to render structure figures.

### Differential Scanning Calorimetry

Protein thermal stability was determined by differential scanning calorimetry (DSC) using a Nano-DSC model 602000 calorimeter (TA instruments). Protein solutions in buffer A (20 mM sodium acetate [pH 5.8], 50 mM NaCl) in the presence and absence of 4 mM PCho were subjected to a temperature increase of 1°C/min from 0°C to 100°C under a pressure of 3 atm, and the evolution of heat was recorded as a differential power between reference (buffer A) and sample (10 μM protein in buffer A) cells. The resulting thermograms (after buffer subtraction) were used to derive thermal transition midpoints (T_m_’s). Fitting to the two-state scaled model provided in NanoAnalyze software was used to obtain a T_m_ value. The experiments were repeated two times with consistent results.

### Isolation of Human Peripheral Blood Mononuclear Cells (PBMCs)

Blood samples were obtained from healthy, consenting donors as Buffy coats (New York Blood Center) and leukopaks (AllCells, Alameda, CA). PBMCs were isolated from peripheral blood by density gradient centrifugation using Ficoll-Paque Plus (GE life sciences).

### Cytolysis Inhibition Assay

Flow cytometry was used to assay permeabilization of the plasma membrane (pore formation) by LukED, PVL and α-toxin in Jurkat cells and primary human immune cells as described previously (Alonzo et al., 2012). Briefly, serial dilutions of C_14_PC were preincubated individually with different concentrations of the LukD and LukF-PV F subunits and α-toxin in V-bottom 96 well-plate for 30 min at 4°C. These mixtures were then added to prestained PBMCs and incubated with the cognate LukE and LukS-PV S partners for 1–1.5 h and with α-toxin for 24 h in a humidified 5% CO_2_ incubator at 37°C. The cytotoxin-treated cells were stained with a viability dye and analyzed by FACS. 50% inhibitory concentration (IC_50_) values are calculated using GraphPad Prism by fitting data to single-slope dose-response curves constrained to 0% and 100% values.

### Staining and FACS Analysis

PBMCs were differentially stained with specific cell surface markers prior to intoxication in order to identify distinct cell populations. Antibodies used for flow cytometric staining included CD3-Alexa 532 (clone UCHT1) (eBioscience, San Diego, CA), CD4-Brilliant Violet 570, CD8-Pacific Blue, CD45RO-APCCy7, CD14-Alexa 700, CD27-PeCy7, CD244 (2B4)-Percp Cy5.5, CXCR1-APC (Biolegend, San Diego, CA), CCR5-PE (BD Biosciences, San Diego, CA) and CCR7-FITC (R&D systems, Minneapolis, MN). After intoxication, cells were collected, washed with phosphate buffered saline and stained with Fixable viability dye eFluor 506 (eBioscience, San Diego, CA). Data were acquired on BD LSRFortessa X-20 instrument (BD Biosciences, CA) using FACSDiva software, iQue Screener PLUS (Intellicyt, MI) using ForeCyt Software or SP6800 Spectral Analyzer (Sony Biotechnology, CA). Data analysis was performed using FlowJo software (TreeStar Inc, Ashland, OR). Statistical analysis was performed using GraphPad Prism 8 software.

### Accession Numbers

Coordinates and structure factors have been deposited in the Protein Data Bank with accession codes 6U33 (LukD), 6U2S (LukD + C_14_PC), 6U3F (LukF-PV + C_14_PC), 6U3T (α-toxin^oH35A^ + C_14_PC), 6U3Y (the PVL heterooctamer + C_14_PC), 6U49 (the α-toxin heptamer + C_14_PC) and 6U4P (the α-toxin^H35A^ heptamer + C_14_PC).

## Acknowledgments

We thank the beamline personnel at the Cornell High Energy Synchrotron Source and the Stanford Synchrotron Radiation Lightsource for data collection, J. Cai for her participation and assistance in the early stage of the project, M. Zhang and Q. Li for technical assistance, and J. Nunberg, N. Kallenbach and J. Lu for comments on the manuscript. This research was supported by NIH grant AI094599.

## Author Contributions

J.L. and M.L. performed the biochemical and biophysical experiments and the co-structure determinations. L.K. and D.U. performed the toxin activity and inhibition measurements. M.L. and D.U. wrote the manuscript with contributions from the other authors. M.L., D.U. and V.T. initiated the project.

